# The *Acinetobacter baumannii* virulence factor Omp33 is important for tolerance to benzalkonium chloride

**DOI:** 10.1101/2023.07.26.550637

**Authors:** Varsha Naidu, Bhumika S. Shah, Karl A. Hassan, Ian T. Paulsen

**Author notes:** **Corresponding author:** School of Natural Sciences, Macquarie University, Sydney, NSW, 2109 Australia.

## Abstract

Benzalkonium chloride (BAC) is widely used in many disinfectant solutions in clinical settings to eradicate potential bacterial pathogens, such as *Acinetobacter baumannii*. We sought to investigate the transcriptomic response of a drug resistant *A. baumannii* isolate, AB5075-UW, on exposure to a sub-inhibitory concentration of BAC. Our transcriptomic analysis found that BAC caused an increase in the expression of genes associated with protein synthesis, such as translation initiation factors, ribosomal proteins and tRNA synthetases. It also induced the expression of genes associated with energy production and central carbon metabolism. We also observed increased expression of peptidoglycan and rod shape determining genes, which may provide increased mechanical strength to withstand osmotic challenges posed by compounds such as BAC. The most highly expressed genes under BAC stress include those that encode the RND efflux pump AdeABC and the *A. baumannii* porin Omp33. Mutants of *adeABC* and its regulator genes *adeRS* had a higher susceptibility to BAC. Disruption of the gene encoding Omp33 also resulted in higher susceptibility to BAC, and complementation of the mutant with *omp33* together with a 450bp upstream region restored tolerance to BAC to parental strain levels (AB5075-UW). Site directed mutagenesis of amino acids associated with Omp33 periplasmic turn (T1), which folds into the lumen of the porin and blocks the channel, suggests that Omp33 may act to prevent entry of BAC into the cell. In previous studies, Omp33 has been described as an important virulence factor in *A. baumannii.* The results presented in this study describe a novel role for Omp33 in BAC tolerance and reveal that *A. baumannii* tolerates BAC stress through a combination of mechanisms.

## Introduction

*Acinetobacter baumannii* has been identified by the World Health Organisation (WHO) as a bacterial pathogen for which new antibiotics are urgently needed (1). *A. baumannii* is of particular concern in health-care settings because of its remarkable ability to survive on abiotic surfaces for extended periods (2). The use of disinfectants in hospitals is an essential measure in controlling the transmission of pathogens such as *A. baumannii* to immunocompromised patients. Surveillance studies comparing resistance to disinfectants in 445 bacterial strains from various genera revealed that *Acinetobacter* were the most resistant species to disinfectants (3). Benzalkonium chloride (BAC) is a quaternary ammonium compound (QAC) used widely in many disinfectant solutions in clinical and community settings to eradicate bacteria. The antibacterial activity of BAC is not well understood, however its amphiphilic nature is thought to cause the general disruption of the bacterial lipid membrane resulting in the loss of osmoregulation and leakage of cellular materials (4). Gram-negative bacteria can be intrinsically more tolerant to antimicrobials than Gram-positives due to their poorly permeable outer membranes and network of active efflux pumps (5). Resistance to QACs has primarily been associated with changes in the bacterial membrane composition in organisms including *Escherichia coli* (6) and *Pseudomonas aeruginosa* (7, 8); reduced expression of outer membrane protein OmpF in *Escherichia coli* (6) and; active extrusion by multidrug efflux pumps including the BcrABC protein in *Listeria monocytogenes* (9), the AdeABC RND efflux system in *A. baumannii* (10) and MexCD-oprJ in *P. aeruginosa* (11). There is growing concern that improper use of disinfectants, such as QACs, could not only lead to increased tolerance to these compounds by bacteria, but promote cross-resistance to antibiotics, due to shared mechanisms of resistance (11–13). The underlying molecular mechanisms by which antibiotic-resistant bacteria such as *A. baumannii* tolerate BAC stress is not well understood. In this study, the global response to BAC was investigated in a multidrug-resistant *A. baumannii* isolate, AB5075-UW using transcriptomics. This enabled the identification of genes in *A. baumannii* that are involved in tolerance to BAC.

## Materials and Methods

### Bacterial strains, growth media and plasmids

The bacterial strains used were *A. baumannii* AB5075-UW, its insertionally-inactivated mutant strains *omp33*171::T26 (AB00516), *adeA*191::T26 (AB05225), *adeB*150::T26 (AB05228), *adeC*102::T26 (AB05231), *adeR*195::T26 (AB05220) and *adeS*132::T26 (AB05215) from the Manoil Laboratory collection (14). Disruption of the *omp33* AB00516 was confirmed by gene-specific primers (Table S1), while disruption of the remaining genes was confirmed previously (15). *E. coli* strain HST08 was used as a host for plasmid propagation. The plasmid used for gene expression was pVRL1Z (16). *E. coli* strains carrying pVRL1Z plasmids were cultured in low-salt LB medium containing 25 μg/ml Zeocin and 300 μg/ml for AB5075-UW strains. Bacterial strains were cultured in cation adjusted Mueller-Hinton broth or agar at 37°C, unless specified otherwise.

### Cell treatments, RNA isolation and Bioinformatic analysis

Three independent *A. baumannii* AB5075-UW cultures were grown overnight in 5ml of Muller-Hinton (MH, Oxoid) broth with shaking (200 RPM) at 37°C. The overnight cultures were re-seeded 1:100 in MH broth and grown for ∼2 hours at 37 °C to exponential phase (OD_600_ = 0.6). Each culture was divided into two culture flasks, one treated with 50 μM benzalkonium chloride, (Sigma-Aldrich) and the other left untreated as a reference. All cultures were grown for an additional 1 hour. Cells were harvested by centrifugation and immediately suspended in Qiazol (Qiagen) to mediate cell lysis.

RNA extraction was carried out using the miRNeasy mini kit (Qiagen) and DNA was eliminated using the TURBO DNA-free kit (Ambion Inc, USA), per the manufacturers’ instructions. The purified total RNA samples were processed and sequenced at the Ramaciotti Centre for Genomics (UNSW). The integrity of total RNA was evaluated using the Agilent Bioanalyzer, followed by rRNA depletion using the Ribo-Zero rRNA removal kit (Gram-negative bacteria) (Illumina, Inc., USA). The cDNA library was generated using the TruSeq® Stranded Total RNA Sample Preparation kit (Illumina, Inc., USA). The samples were sequenced on an Illumina HiSeq4000 platform yielding approximately 20 million 100 bp paired end (PE) reads per sample.

The reads were analysed using EDGE-pro (Estimated Degree of Gene Expression in Prokaryotic Genomes) (17), where reads were mapped to the *A. baumannii* AB5075-UW reference genome (Genbank accession: CP008706.1) using Bowtie2 and raw read counts were generated. The count data was analysed with the R package DESeq2 to identify genes with differential expression (18).

### PCR, Gene cloning and complementation

The coding region of the Omp33 gene with an upstream region of 450 bp was amplified by PCR using gene-specific primers (Table S1) with AccuPrime™ Pfx SuperMix (Invitrogen) and cloned into the pVRL1Z plasmid using restriction enzymes *EcoR*I and *Not*I. Insertion of the genetic construct into the plasmid was confirmed via PCR, followed by Sanger sequencing using universal M13 forward and reverse primers (Table S1). Recombinant plasmids were transformed and recovered from an *E. coli* HST08 strain using the QIAprep Spin Miniprep Kit (Qiagen). Recombinant and empty pVRL1Z plasmids were transformed into electro-competent *A. baumannii* strains as previously described (19).

### Substitution of amino acid residues via site-directed mutagenesis (SDM)

The substitution of amino acids via SDM was carried out using the QuikChange II Site-Directed Mutagenesis Kit (Agilent Technologies) per the manufacturer’s instructions. Briefly, the recombinant pVRL1Z plasmid carrying the Omp33 gene and 450bp upstream was transformed into and recovered from the *E. coli* DH5α strain. Mutant strand synthesis was performed as per manufacturer’s instructions using codon specific mutagenic primers (Table S1). The parental (nonmutated) plasmid was digested with *Dpn I* followed by transformation in XL1-Blue supercompetent cells. Mutated plasmids were recovered using the QIAprep Spin Miniprep Kit (Qiagen) and sent for Sanger sequencing using universal M13 forward and reverse primers (Table S1). Sequence verified mutated plasmids were transformed into electrocompetent AB00516 (Δ*omp33*) strains as previously described (19).

### Antimicrobial Susceptibility Testing

Susceptibility assays were carried out with stationary phase cultures in Mueller-Hinton broth using the micro broth dilution method as described (20). Cultures were incubated at 37°C with shaking at 200 RPM and cell growth was determined by measuring the absorbance (OD600) as 24h endpoint measurements in the PHERAstar FSX (BMG LABTECH, Germany). The growth rate of strains encoding Omp33 site directed mutants was measured in the BioTek LogPhase 600 Microbiology Reader (Agilent) continuously every 10 mins for 24h at 37°C with shaking at 500 RPM.

## Results and Discussion

### Global transcriptional response to benzalkonium chloride (BAC)

An international clonal complex I *A. baumannii* strain, AB5075-UW (21), with a benzalkonium chloride minimum inhibitory concentration (MIC) of 100 μM (Figure 1) was exposed to a sub-inhibitory concentration (50 μM) of BAC for 1h. Subsequently, the transcriptional response was analysed via RNA-Sequencing, providing quantitative expression data for 3,983 (out of 3987) genes (14). In total, 1041 genes showed significantly higher transcript levels, while 1046 genes had significantly lower transcript levels (p-adjusted, < 0.05) when exposed to BAC. Functional class analysis of these genes assigned them into four core Clusters of Orthologous Group (COG) categories: cellular processes and signalling (11.88%); information storage and processing (17.72%); metabolism (33.78%) and poorly characterised (36.6%).

**Figure 1.**
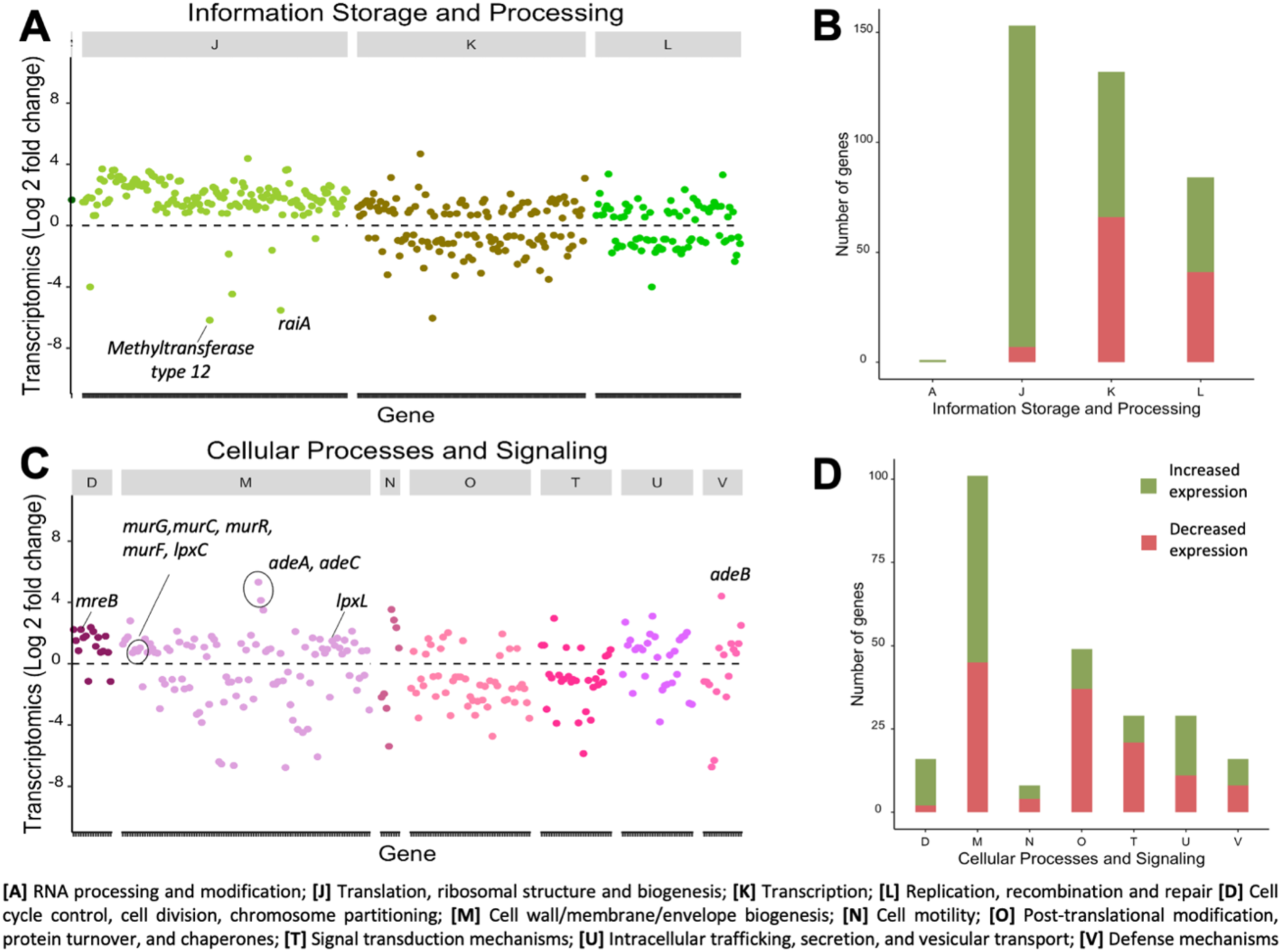
**(A)** Functional class analysis of genes based on COG category ‘Information, Storage and Processing’. Each point in the graph represents a single ORF according to its position within the genome, arranged according to their COG and their fold-change (log_2_) in expression on the y-axis. **(B)** Number of genes in the COG category ‘Information, Storage and Processing’ that had either increased (green) or decreased (orange) expression (p-adjusted <0.05). **(C)** Functional class analysis of genes based on COG category ‘Cellular Processing and Signalling’. **(D)** The number of genes in the COG category ‘Cellular Processing and Signalling’ that had either increased (green) or decreased (orange) expression (p-adjusted <0.05).

#### Protein Synthesis

A total of 146 out of 153 genes under the category “translation, ribosome structure and biogenesis”, associated with protein translation had increased transcript levels in response to BAC. These include tRNA synthetase genes, genes encoding translation initiation factors, as well as the *S10*, *spc* and *alpha* gene clusters that encode ribosomal proteins (Figures 2A and 2B). In contrast, the protein factor RaiA encoded by the gene *yfiA* (22, 23), a translation inhibitor, had decreased expression (Figure 2A). Tetracycline and other antibiotics exert their antibacterial activity by inhibiting protein synthesis in bacteria. An increase in the expression of these genes has been observed in response to tetracycline (our unpublished data) and tigecycline stress (24) in *A. baumannii*. Furthermore, adaptive evolution experiments with BAC in *E. coli* have shown increased expression of RpsF protein (30S ribosomal subunit S6) in cells that evolved to have a high tolerance for BAC (25). This suggests that genes associated with translation may play a role in BAC tolerance and raises the question of whether BAC may target protein translation in addition to its direct effects on membrane integrity.

**Figure 2.**
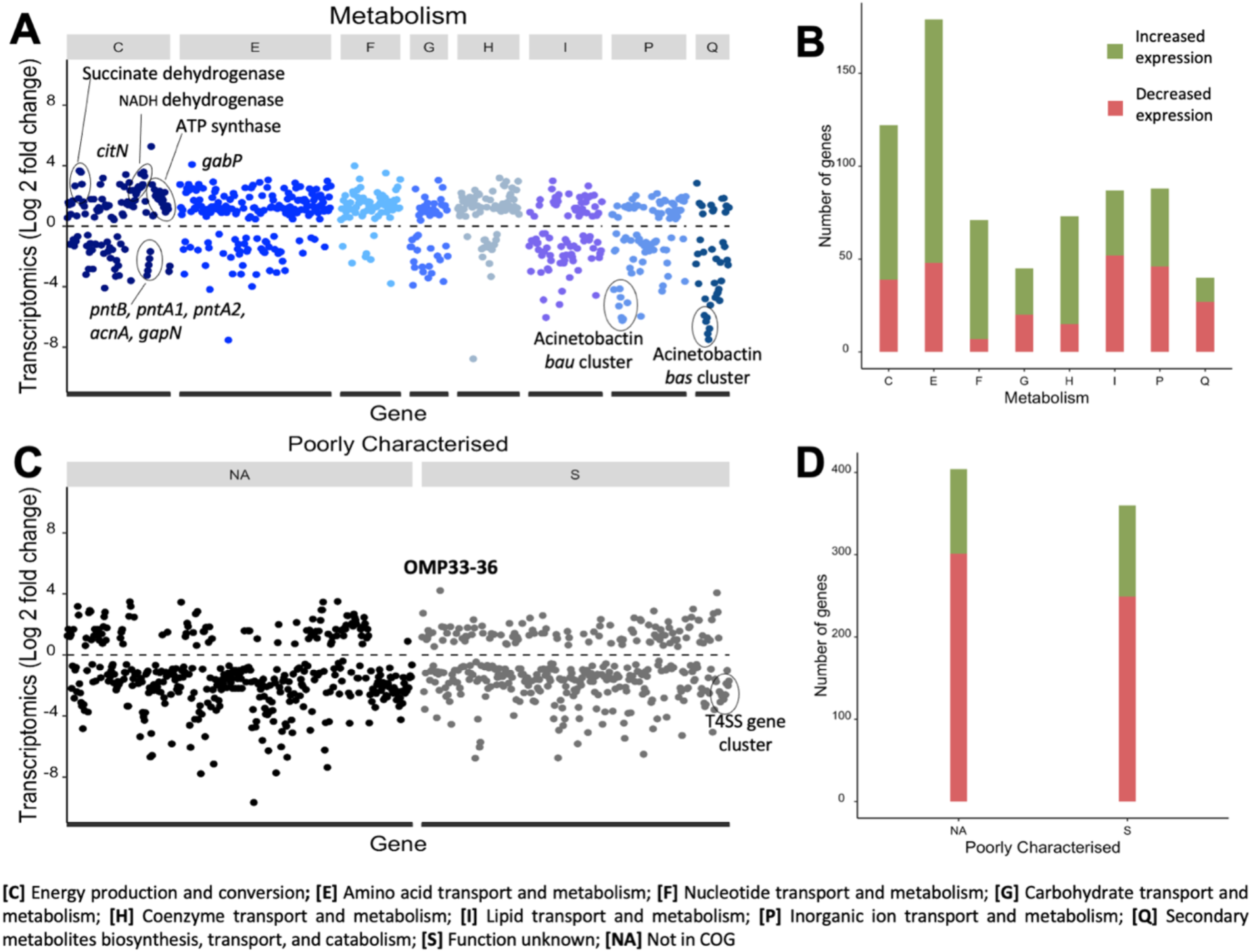
**(A)** Functional class analysis of genes based on COG category ‘Metabolism’. Each point in the graph represents a single ORF within the genome, arranged according to their COG and fold-change (log_2_) in the expression on the y-axis. **(B)** Number of genes in the COG category ‘Metabolism’ that had either increased (green) or decreased (orange) expression (padj <0.05). **(C)** Functional class analysis of genes based on COG category ‘Poorly characterised’. **(D)** The number of genes in the COG category ‘Poorly characterised’ that had either increased (green) or decreased (orange) expression (padj <0.05).

#### Cell wall, cell shape and Lipid A biosynthesis

Peptidoglycan (PG) biosynthesis genes *murA, murB, murC, murD, murE, murF* and *murG* had increased expression in *A. baumannii* strain AB5075-UW following exposure to BAC (Figure 2C). PG is an important cell wall component that maintains cell shape and provides mechanical strength to withstand osmotic challenges (26). Genes associated with cell shape determination including *mreB, mreC* and *mreD* also had increased expression in response to BAC stress (Figure 2C). The MreB protein has been shown to play an essential role in maintaining the rod shape in many bacteria, regulating peptidoglycan synthesis (27) and exerting mechanical stiffness in cells by tethering to the cell wall (28). In addition, genes associated with Lipid A biosynthesis including *lpxA, lpxB, lpxC, lpxH, lpxK* and *lpxL* had increased expression (Figure 2C). Lipid A anchors the lipooligosaccharide (LOS) to the cell’s outer membrane (29). The increased expression of genes involved in cell envelope maintenance, suggests that *A. baumannii* may rely on such external features to confer tolerance to BAC, or is a response to cell envelope damage caused by BAC.

#### Cellular respiration and metabolism

Genes encoding enzymes involved in respiration, including the F-type ATP synthase, NADH dehydrogenase I, cytochrome O ubiquinol oxidase and succinate dehydrogenase, had increased expression under BAC stress (Figure 3A). Recently BAC and other biocides have been shown to collapse membrane potential at sub-inhibitory concentrations (15). Therefore, these transcriptional changes are most likely a response to a loss in membrane potential. Our transcriptomic work also shows increased expression of genes associated with amino acid biosynthesis, nucleotide metabolism and tricarboxylic acid (TCA) cycle, possibly to try to compensate for a drop in membrane potential or alternatively an accelerated metabolism to meet higher energy demands for processes such as translation and transcription.

**Figure 3.**
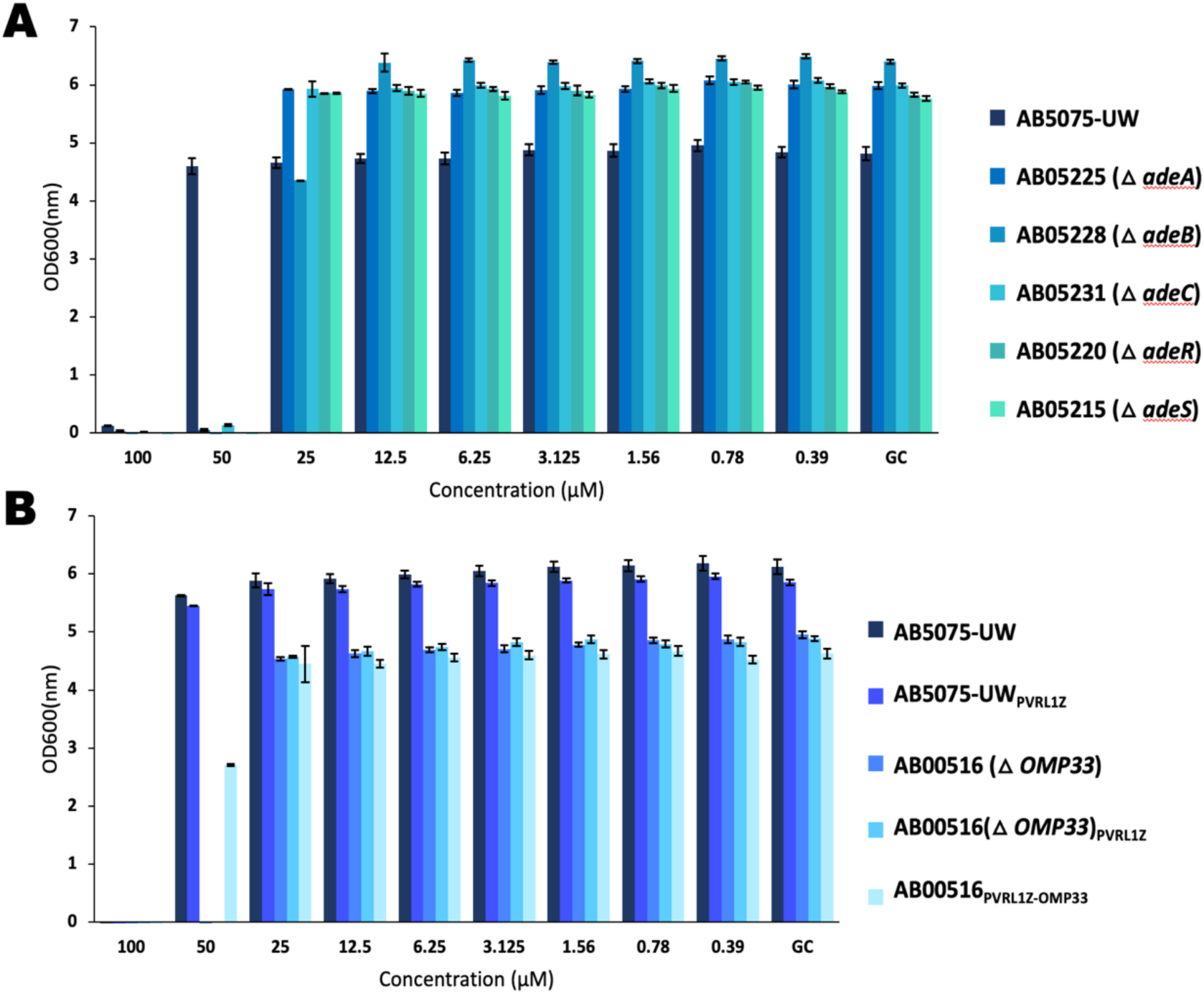
The susceptibility of AB5075-UW, and transposon insertion mutants were tested using the broth microdilution method. Each strain was subjected to either different concentrations of benzalkonium chloride or no benzalkonium chloride. The optical density was measured after 24h as an endpoint measurement. Data bars represent the geometric mean ± SE (*n* = 8) of two biological replicates. **(A)** Strains Δ*adeA* (AB05225), Δ*adeB* (AB05228), Δ*adeC* (AB05231), Δ*adeR* (AB05220) and Δ*adeS* (AB05215) were more susceptible to benzalkonium chloride when compared to parental strain (AB5075-UW). **(B)** Growth of strains AB00516 (ΔOmp33), and AB00516 expressing empty pVRL1Z plasmid was significantly inhibited in the presence of 50 μM of benzalkonium chloride. Complementing AB00516 (Δ Omp33), with the Omp33 plus upstream region of 450 bp (AB00516_pVRL1Z-Omp33_) restored partial susceptibility to 50 μM of benzalkonium chloride.

#### Cell Signalling

Gene clusters with lower relative transcript abundance under BAC stress include the acinetobactin gene cluster (Figure 3A), composed of 18 genes. The acinetobactin cluster comprises acinetobactin biosynthesis, utilization and release genes. Acinetobactin is a siderophore that plays a vital role in acquiring iron during iron limitation conditions and in virulence (30). Genes within the F-like type IV secretion system (T4SS) cluster encoded in the AB5075-UW plasmid, p1, also showed reduced transcript abundance after BAC exposure (Figure 3C). The T4SS is important for conjugative transfer of DNA, plasmids, and other mobile genetic elements (31).

#### Transporters and Porins

Transcripts encoding several transport proteins were more abundant after BAC exposure, including the AdeABC RND efflux system (Figure 2C), GABA permease (GabP) (Figure 3A), uracil-xanthine permease and the CitN citrate transporter (Figure 3A). The GABA permease, which is driven by membrane potential, is responsible for the uptake of gamma-aminobutyric acid (GABA) (32). In *E. coli*, the transport of GABA is dependent on the presence of phosphatidylethanolamine (PE) within the cell membrane (33) and it is generated intracellularly from glutamic acid as a glutamate-dependent acid stress response (34). The increased expression of *gabP* and GABA synthesising genes *gabD* and *gabT* in AB5075-UW may indicate a similar stress response role in BAC tolerance.

Overexpression of AdeABC has been shown to provide resistance to BAC (10). The expression of the AdeABC efflux genes is regulated by the two-component transcriptional regulatory system AdeRS where AdeS is a sensor kinase and AdeR is the cognate response regulator (35, 36). The expression of *adeRS* was not changed in the presence of BAC. The role of these genes in conferring tolerance to BAC was further investigated by MIC tests using the Δ*adeA,* Δ*adeB,* Δ*adeC,* Δ*adeR and* Δ*adeS* transposon mutants in AB5075-UW (14). The *adeABC* and *adeRS* transposon mutants had a two-fold increase in susceptibility to BAC compared to the parental strain (Figure 4), suggesting an important role in tolerance.

**Figure 4.**
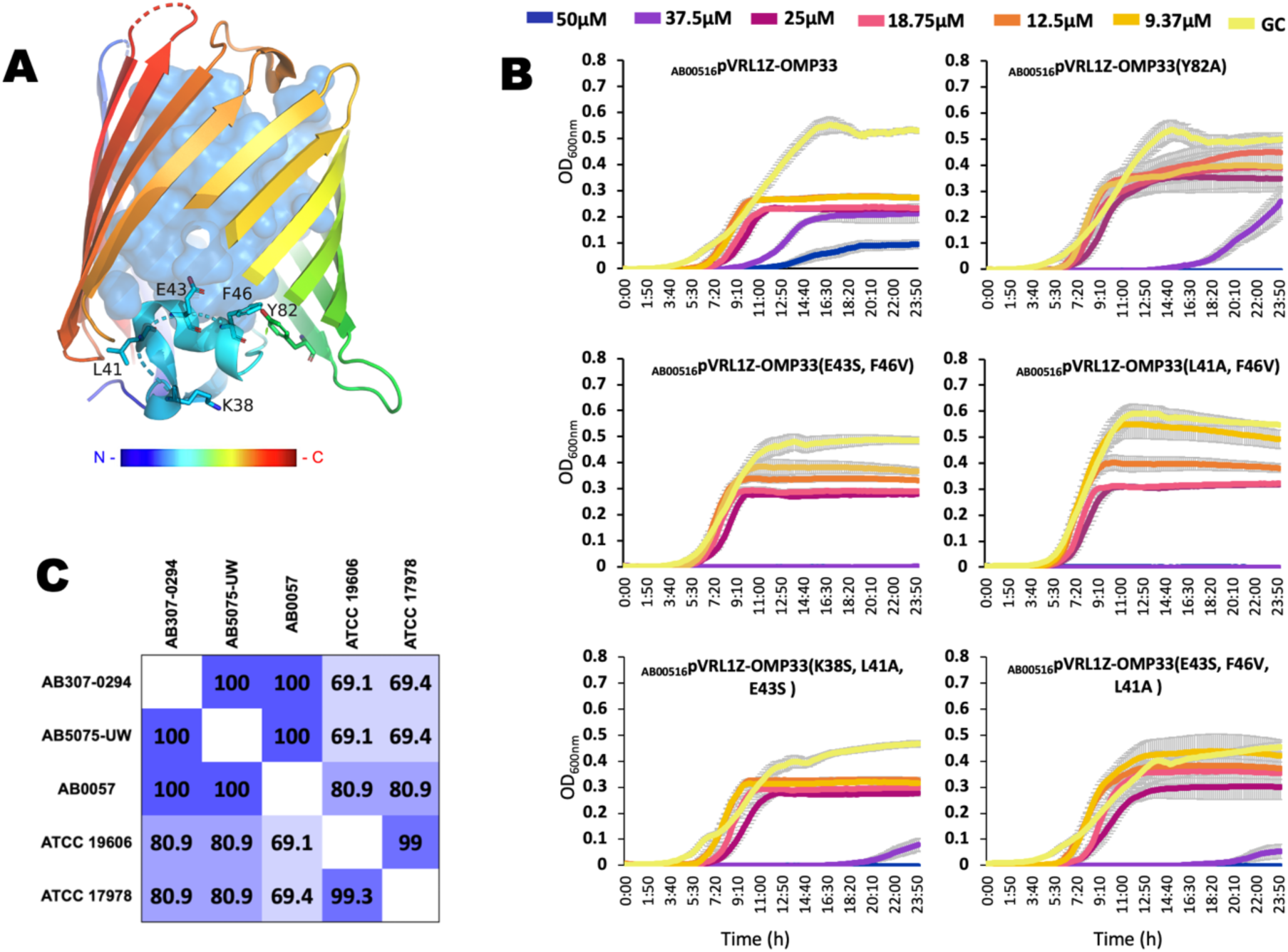
(A) Solved structure of Omp33 (PDB – 6GIE) from the *A. baumannii* strain AB307-0294 (38) depicting residues that were mutated via site directed mutagenesis in the Omp33 periplasmic turn T1. Residues L41A, E43S, F46V, Y82A, and K38S were introduced as different combinations in Omp33, resulting in five different Omp33 mutants (panel B). (B) Plasmid pVRL1Z carrying the Omp33 gene with the different mutation combinations were transformed into the AB00516 (ΔOmp33) strain and their susceptibility to different concentrations of benzalkonium chloride was measured as optical density (OD600nm) over a 24h period. The growth assays represent three independent experiments. Each line represents geometric mean ± SE (*n* = 12) of three biological and four technical replicates. (C) Pairwise amino acid sequence similarity scores for Omp33 proteins from *A. baumannii* sequence type I (AB307-0294, AB5075-UW and AB0057); sequence type 2 (ATCC 17978) and sequence type 6 (ATCC 19606). These values were determined using MatGat version 2.0 with a BOLSUM 50 scoring matrix (43).

The transcriptomic data reveal changes in the expression of genes encoding several porins including CarO, ABUW_0166, oprE3 and Omp33 (also known as Omp34 or MapA)(37). While genes encoding CarO and ABUW_0166 showed reduced transcript abundance (-3.97 and -4.12 log 2-fold change, respectively) and oprE3 showed a moderate increase in expression (2.28 log 2-fold change), the gene encoding Omp33 showed a significant increase in expression (4.21 log 2-fold change) following BAC exposure.

### Porin Omp33 plays an important role in tolerance to benzalkonium chloride

To investigate whether Omp33 provides tolerance to BAC, we performed MIC tests with the Omp33 transposon mutant strain (AB00516) (14, 20). We found that the AB00516 (Δ*omp33*) strain was two-fold more susceptible to BAC than the parental strain, AB5075-UW (Figure 3). The gene encoding Omp33 together with its predicted endogenous promoter and operator sequence (450bp upstream region of Omp33) was cloned into the pVRL1Z vector (16), and the resulting plasmid pVRL1Z_Omp33_, was used to complement the AB00516 (Δ*omp33*) strain (Figure 3). MIC tests were performed on AB00516 (Δ*omp33*) complemented with pVRL1Z_Omp33_ and empty vector controls AB5075-UW_pVRL1Z_ and AB00516_pVRL1Z_. We found that complementing AB00516 mutant strain with pVRL1Z_Omp33_ restored partial tolerance to BAC.

The published x-ray crystal structure of Omp33 suggests that it is a 14-β stranded barrel, with extended periplasmic turns which fold into the lumen of the pore and block the channel completely, acting as a gated channel (38). We sought to investigate the role of this gated channel by generating five different combinations of amino acid substitutions associated with the periplasmic turn (T1), via site directed mutagenesis (Figure 4A). This resulted in five different *omp*33 genes with unique combination of mutations. The mutated *omp*33 genes with the 450bp upstream region was cloned into the shuttle vector pVRL1Z. MIC tests were performed with all five mutant Omp33 genes, and AB00516_pVRL1Z-OMP33_ as control (Figure 6B). We found that all five mutated Omp33 proteins, potentially in an open conformation, were more susceptible to BAC in comparison to the parental strain and the AB00516 mutant strain with the recombinant plasmid (pVRL1Z_Omp33_) (Figure 4B).

Omp33 is one of the most important virulence factors in *A. baumannii*, which is associated with the adherence and subsequent invasion of human epithelial cells (39) and is considered an important vaccine target against *A. baumannii* infections (40). Omp33 has been shown to induce apoptosis and modulate autophagy, allowing *A. baumannii* to persist inside autophagosomes (41). In addition to its role in virulence, Omp33 has also been associated with carbapenem resistance, whereby its decreased expression has been associated with reduced susceptibility to carbapenems (37). Omp33 is found on the surface of the bacterium and acts as a channel for the passage of water and the protein itself can be released to the environment in outer membrane vesicles (OMVs) (41).

Sequence conservation analysis of Omp33 in *Acinetobacter* species shows that this protein can be divided into three groups, with an average amino acid polymorphism greater than 80% (42). Of these three groups, group III is the least conserved (42). Omp33 from AB5075-UW belongs to group III. Sequence similarity analysis using MatGat and MUSCLE shows that the Omp33 sequences from *A. baumannii* strains AB5075-UW, ATCC19606 and AB307-0294 are more closely related than those from ATCC17978 and AB0057 (Figure 4C and S1).

The data presented in this study for the first time demonstrates that Omp33 is important or tolerance to benzalkonium chloride in the *A. baumannii* strain AB5075-UW. The phenotypic characteristics of Omp33 uncovered in this study, suggest that the Omp33 gate may play a role in tolerance to BAC. Omp33 may provide tolerance by either stabilising the osmotic pressure or by helping to maintain membrane structural integrity. The identification of Omp33 role in BAC tolerance demonstrates the significant value of genome-wide expression studies to identify novel antimicrobial targets and tolerance factors in multi-drug resistant bacteria. Furthermore, the use of benzalkonium chloride could provide selective pressure for maintenance of Omp33 virulence factor.

## Acknowledgement

This work was supported by the NHMRC (National Health and Medical Research Council) project grants APP1120298 to ITP and KAH. ITP is supported by ARC (Australian Research Council) Laureate Fellowship FL140100021. KAH is supported by an ARC Future Fellowship FT180100123.

## Transparency declaration

None to declare.

---

**Figure S1.**
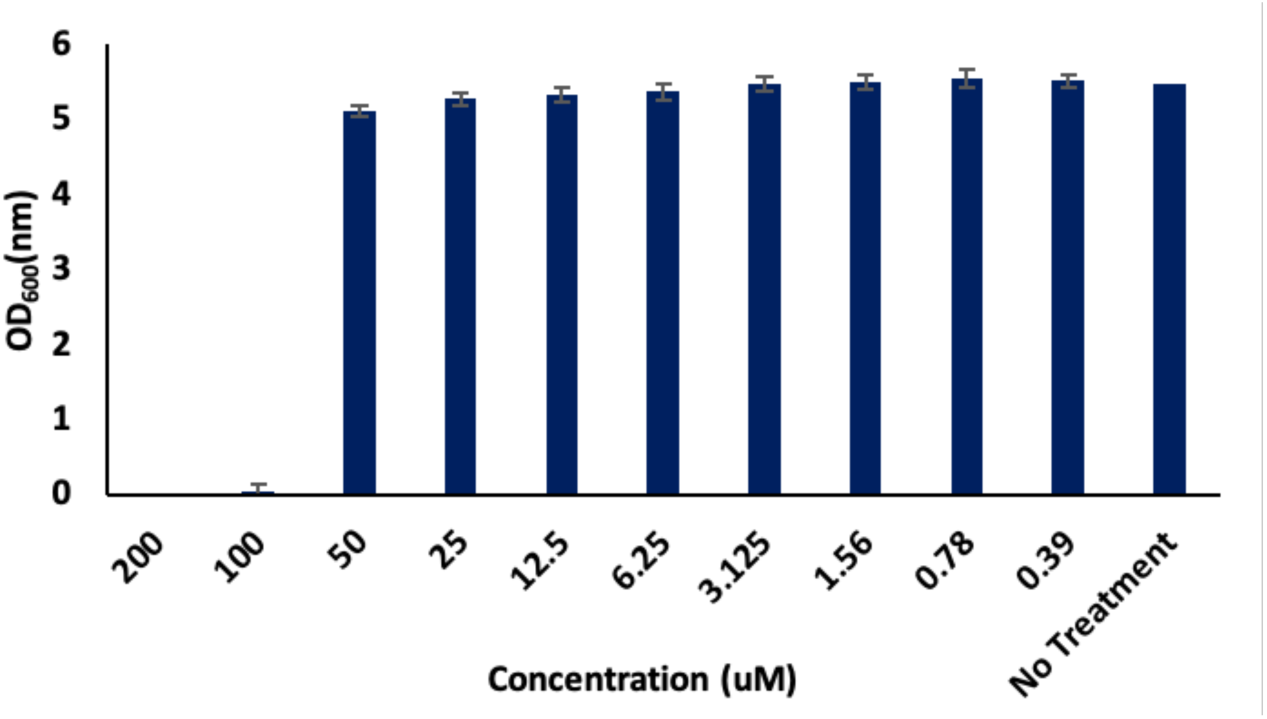
The susceptibility of AB5075-UW to benzalkonium chloride was determined using a minimum inhibitory concentration (MIC) broth microdilution method. The MIC was determined to be 100 μM.

**Figure S2.**
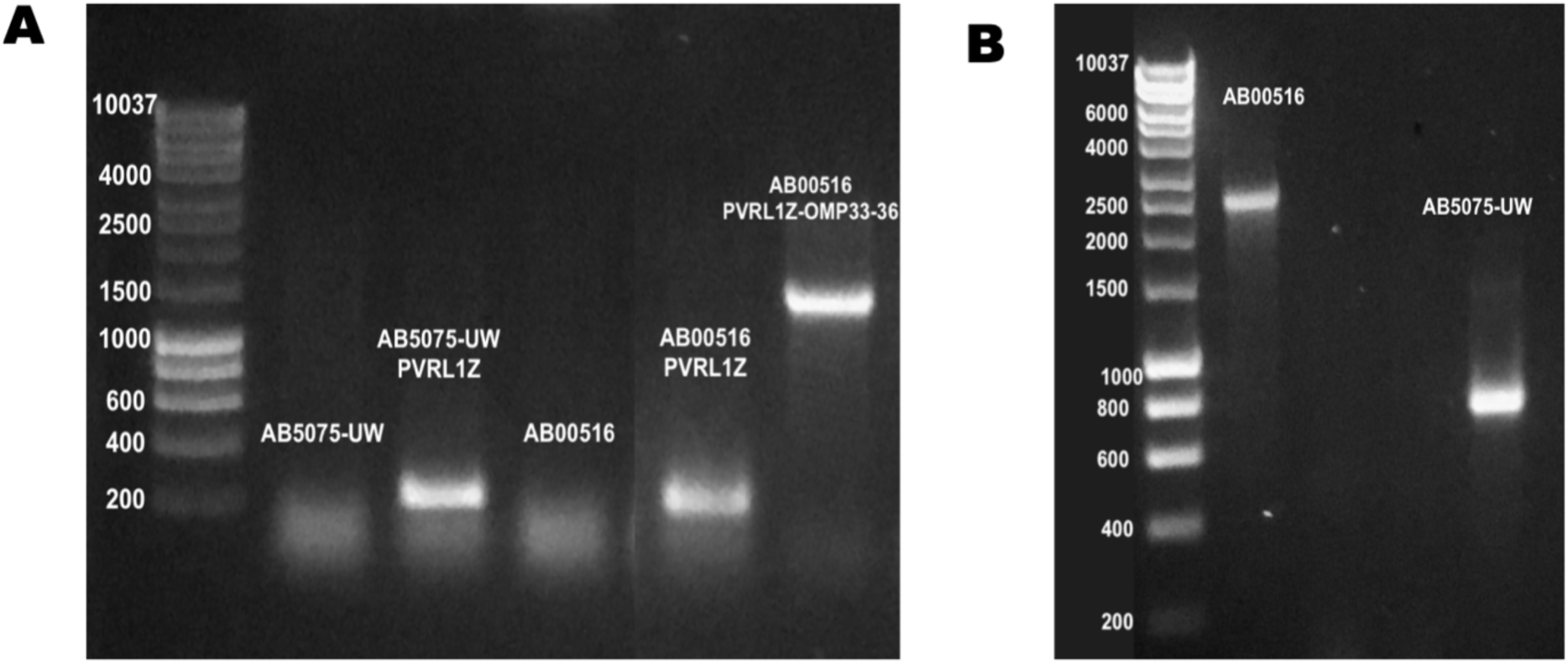
**(A)** M13 universal primers were used to confirm the insertion of Omp33 gene with a 450bp upstream region. The sizes of the molecular weight standards are shown on the left (lane 1). Primers used for the reaction correspond to AB5075-UW (Lane 2), AB5075-UW expressing empty vector -pVRL1Z (Lane 3), AB00516 (Lane 4), AB00516 expressing empty vector -pVRL1Z (Lane 5) and AB00516 expressing Omp33 gene with its presumably endogenous promoter (450bp upstream region of Omp33) on a pVRL1Z plasmid (lane 6). **(B)** *Omp33* gene-specific primers were used to confirm insertional inactivation of *Omp33* gene. The sizes of the molecular weight standards are shown on the left (lane 1). Lane 2 corresponds to strain AB00516 (*Omp33* gene disrupted by the T26 transposon). Lane 3 corresponds to AB5075-UW (Wildtype) strain.

**Table S1.**
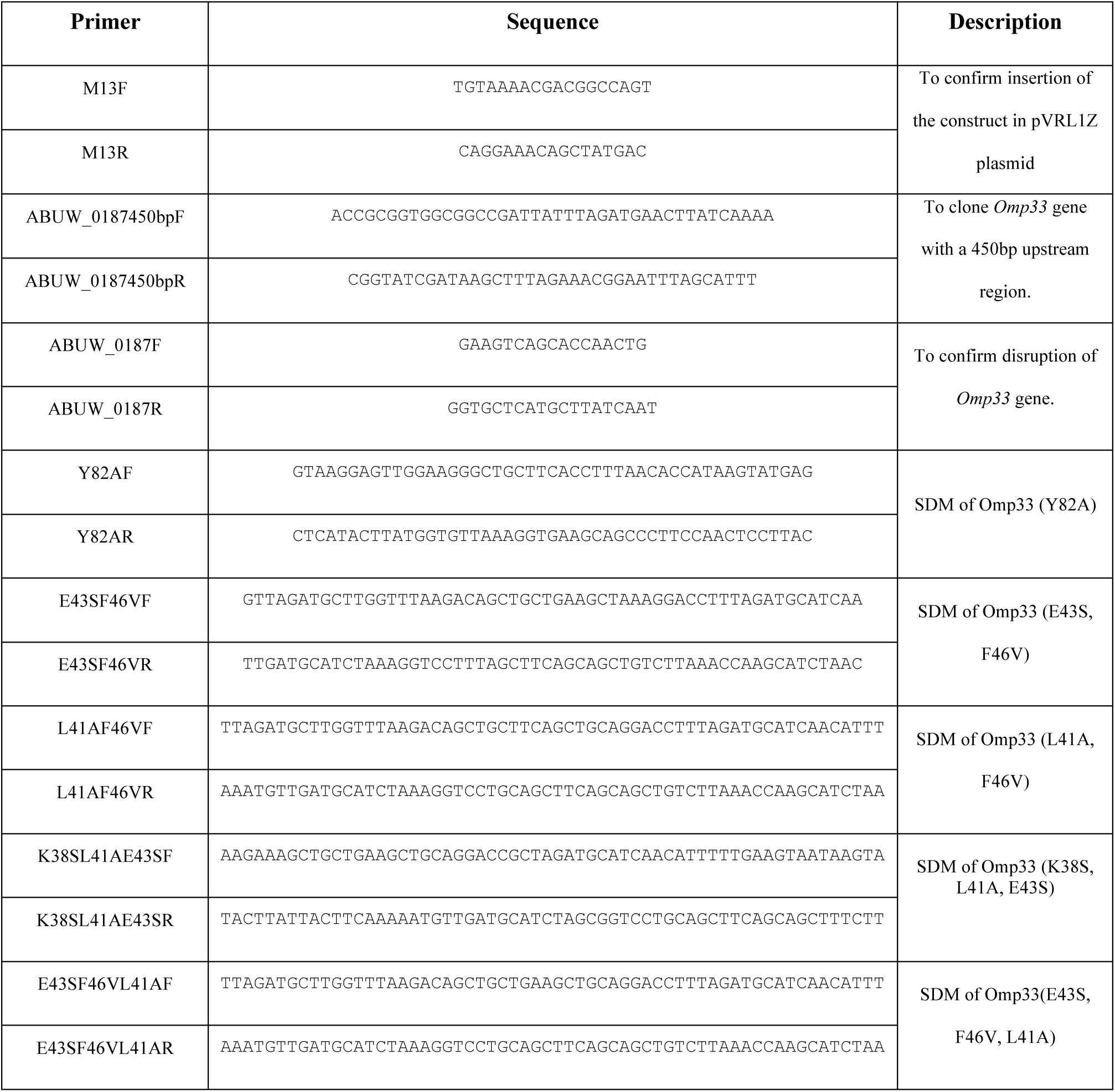
Primers used in this study.

**Figure S3.**
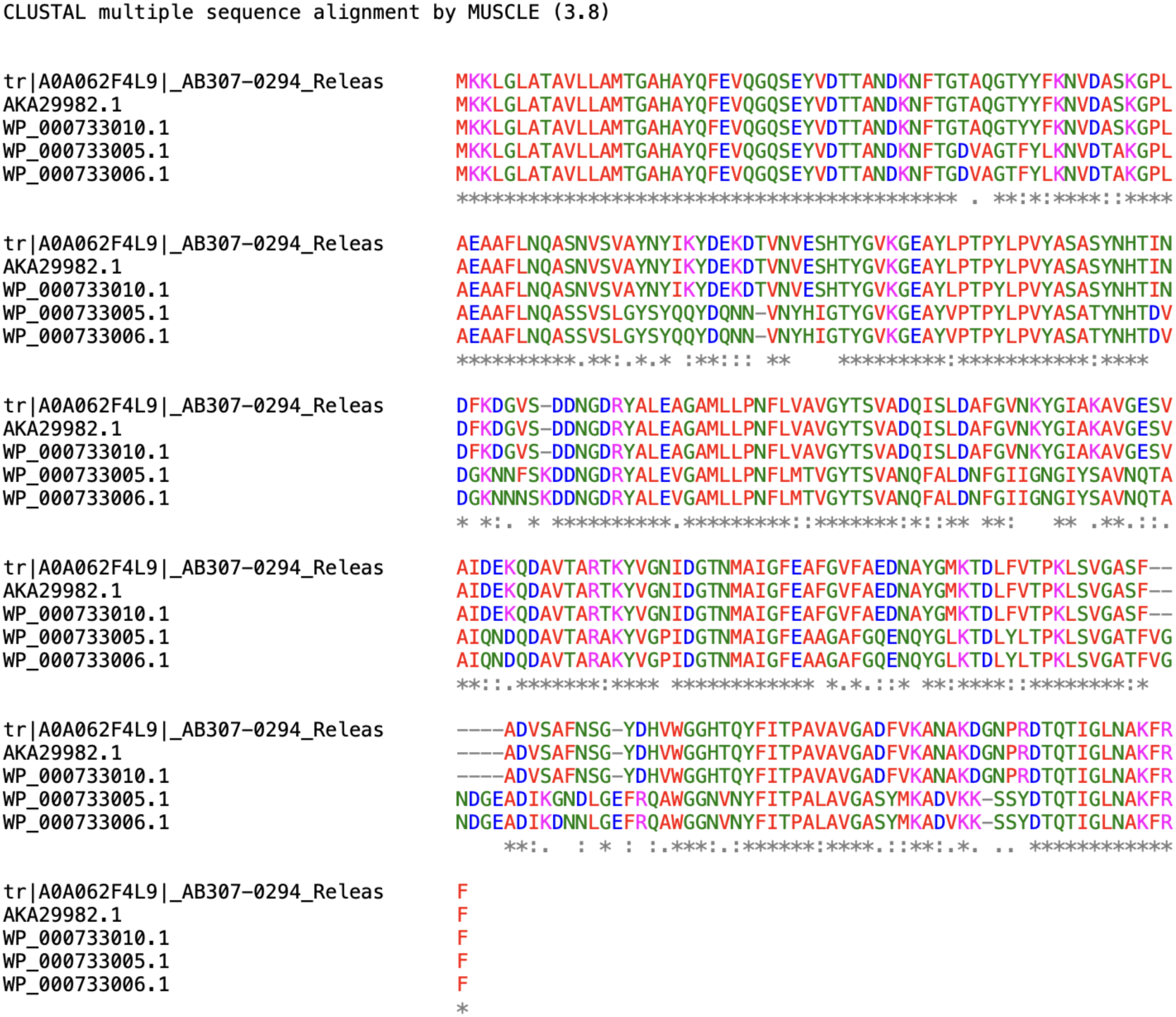
Multiple sequence alignment of the Omp33 amino acid sequence from *A. baumannii* strains AB307-0294 (tr|A0A062F4L9), AB5075-UW (AKA29982.1), ATCC19606 (WP_000733010.1), ATCC17978 (WP_000733005.1) and AB0057 (WP_000733006.1) using MUSCLE. Omp33 sequences from AB5075-UW, ATCC19606 and AB307-0294 show a higher similarity in comparison to sequences from ATCC17978 and AB0057.

## Notes

### Competing Interest Statement

The authors have declared no competing interest.

